# Temporal patterns of haplotypic and allelic diversity reflect the changing selection landscape of the malaria parasite *Plasmodium falciparum*

**DOI:** 10.1101/2024.06.23.600231

**Authors:** Angela M. Early, Stéphane Pelleau, Lise Musset, Daniel E. Neafsey

## Abstract

Populations of the malaria parasite *Plasmodium falciparum* regularly confront orchestrated changes in frontline drug treatment that drastically alter the parasite’s selection landscape. When this has occurred, the parasite has successfully adapted to the new drugs through novel resistance mutations. These novel mutations, however, may emerge in a genetic background already shaped by prior drug selection. In some instances, selection imposed by distinct drugs has targeted the same loci in either synergistic or antagonistic ways, resulting in genomic signatures that can be hard to attribute to a specific agent. Here, we use two approaches for detecting sequential bouts of drug adaptation: haplotype-based selection testing and temporal changes in allele frequencies. Using a set of longitudinally acquired samples from French Guiana, we determine that since the introduction of the drug artemether-lumefantrine (AL) in 2007 there have been rapid hard selective sweeps at both known and novel loci. We additionally identify genomic regions where selection acted in opposing directions before and after widespread AL introduction. At four high-profile genes with demonstrated involvement in drug resistance (*crt*, *mdr1*, *aat1*, and *gch1*), we saw strong selection before and after drug regime change; however, selection favored different haplotypes in the two time periods. Similarly, the allele frequency analysis identified coding variants whose frequency trajectory changed sign under the new drug pressure. These selected alleles were enriched for genes implicated in artemisinin and/or partner drug resistance in other global populations. Overall, these results suggest that drug resistance in *P. falciparum* is governed by known alleles of large effect along with a polygenic architecture of more subtle variants, any of which can experience fitness reversals under distinct drug regimes.

## INTRODUCTION

Humans have created a complex selection landscape of multiple drug pressures to which pathogens continuously adapt (McClure and Day 2014; Huijben and Paaijmans 2017). For pathogens like the malaria parasite *Plasmodium falciparum*, drug use is generally regulated at a national level, with frontline drugs being replaced when clinical resistance surpasses a predefined threshold (WHO 2023). Overall, this translates into parasite populations experiencing sequential bouts of relatively uniform long-term selection pressure. With the introduction of each new drug (or drug combination), *P. falciparum* has successfully adapted with novel resistance mutations arising in at least some global populations (Anderson and Roper 2005; Blasco et al. 2017). Often, the mechanisms of resistance to distinct drugs are complex and overlapping. Pleiotropic effects of these mutations may include protection against related drugs, enhanced susceptibility to other drugs (Shafik et al. 2022), or fitness tradeoffs when no drug is present (Rosenthal 2013). Selection caused by distinct drugs may therefore target the same region of the genome in either synergistic or antagonistic ways.

Selection at the *P. falciparum chloroquine resistance transporter* (*crt*) locus illustrates these potentially complex evolutionary dynamics. Chloroquine resistance in *P. falciparum* was initially mapped to mutations in this membrane transport protein (Su et al. 1997; Fidock et al. 2000), most prominently the mutation CRT^K76T^ (Djimdé Abdoulaye et al. 2001; Sidhu et al. 2002; Ecker et al. 2012; Kim et al. 2019). Subsequent studies found that CRT variants also impact resistance to additional drugs including other 4-Aminoquinolines (such as amodiaquine and piperaquine) (Sá et al. 2009; Beshir et al. 2010; Gabryszewski et al. 2016) and Arylaminoalcohols (quinine, mefloquine, halofantrine, and lumefantrine) (Bellanca et al. 2014). In several known instances, individual alleles or specific allelic combinations show collateral sensitivity, enhancing resistance to one drug class while increasing susceptibility to another (Sidhu et al. 2002; Sisowath et al. 2009; Bellanca et al. 2014; Veiga et al. 2016; Ross et al. 2018; Wicht et al. 2020). CRT^K76T^ carries a fitness cost in the absence of drug (Liu et al. 1995; Ord et al. 2007), and so far, two distinct evolutionary paths have been observed at the locus when chloroquine pressure is removed in natural populations: (1) a resurgence of the wildtype allele CRT^K76^ (*eg*, Malawi (Kublin et al. 2003; Laufer et al. 2010), Kenya (Mwai et al. 2009), Hainan (China) (Wang et al. 2005), and Cameroon (Ndam et al. 2017)) and (2) the acquisition of new CRT mutations that mitigate the fitness cost of CRT^K76T^ or provide resistance to a newly introduced drug (*eg,* CRT^C350R^ in the Guiana Shield (Pelleau et al. 2015; Florimond et al. 2024) and the Cam734 CRT allele in Southeast Asia (Durrand et al. 2004; Petersen et al. 2015)).

The generalizability of these CRT patterns to other loci is unclear as the selection dynamics at other resistance genes are not as thoroughly studied. However, we highlight here one similarity: As with *crt*, surveys in South America and elsewhere have found resistance alleles persisting in *dhfr* and *dhps*, genes involved in antifolate resistance, after these drugs had been officially replaced (McCollum et al. 2007; Guerra et al. 2022). It is unknown what maintains these alleles in the population despite their documented *in vitro* fitness costs in the absence of drug, but evidence suggests that, like with CRT, new mutations in these genes may enable the persistence of older variants by compensating for their fitness cost after the drug is removed (Brown et al. 2010; Salas et al. 2023). Compensatory mutations may also happen in *trans* (Jiang et al. 2008); the most widely studied interaction is between CRT and MDR1, isoforms of which contribute to drug resistance in non-additive ways (Sá et al. 2009; Wicht et al. 2020). In summary, there is growing evidence that the evolution of *P. falciparum* drug resistance is shaped by both pleiotropic effects and compensatory mutations, suggesting that distinct drug regimes may share common genomic targets to a greater extent than is currently recognized. The genomic signatures left by sequential sweeps at the same loci may go undetected with standard methods and require new analysis approaches.

Here, we more broadly assess the effects and targets of recurrent bouts of drug selection with whole-genome data from a *P. falciparum* population in French Guiana, a low transmission region of South America (Musset et al. 2014). We have chosen to focus on a low transmission region because the parasite shows different, somewhat counterintuitive evolutionary dynamics in small versus large populations: despite the greater effect of drift, small populations establish novel adaptive alleles faster than large populations (Anderson and Roper 2005; McClure and Day 2014; Menard and Dondorp 2017). We use longitudinal data that spans the implementation of novel malaria measures throughout the Guiana Shield, which included the introduction of artemisinin combination therapy (ACT) as the new frontline drug treatment. In French Guiana specifically, these interventions decreased malaria incidence by a factor of 15, and as we previously reported, *P. falciparum* relatedness measurably increased across the time span (Early et al. 2022). At the same time, selection remained efficacious as evidenced by measurable signatures of purifying selection, which makes selection testing feasible in the population (Early et al. 2022).

We conduct haplotype-based selection scans at time points before and after the change in drug regime to disentangle the temporal order of selection events and assess the likelihood of hard versus soft selective sweeps under distinct drug regimes. At both time periods, we find evidence of sweeps at known resistance-conferring genes and at novel loci that may represent region-specific modes of adaptation. Next, we identify genes that fit a pattern of selection pressure reversal by analyzing haplotype changes and allele frequency trajectories through time. We find this evolutionary pattern at dozens of resistance-implicated loci accompanied by both hard and soft sweep signatures. This suggests that fitness trade-offs manifest genome-wide and are not limited to large-effect genes. Overall, these results show that selection under distinct drug regimes recurrently targets the same loci, highlighting the importance of considering the identity not just the location of selected haplotypes.

## RESULTS

### Change in frontline drug did not reverse the high prevalence of known large-effect drug resistance alleles in French Guiana

French Guiana is located within the Guiana Shield region of South America and harbors a low *P. falciparum* endemicity that is concentrated in remote regions, particularly sites connected with gold mining (Musset et al. 2014; Douine et al. 2020). Movement of people and parasites throughout the Guiana Shield is well documented (Douine et al. 2017; Mathieu et al. 2021) and previous genetic clustering analyses of South American *P. falciparum* have shown that the Guiana Shield region harbors a genetically distinct parasite population (Yalcindag et al. 2012; Carrasquilla et al. 2022; Lefebvre et al. 2023). *P. falciparum* gene flow therefore occurs across a region that is exposed to the varied public health measures in Brazil, French Guiana, Suriname, Guyana, and Venezuela (Sanna et al. 2024). Across our sampling period, drug use varied throughout these countries, with a general trend of artemisinin combination therapy (ACT) selection pressure intensifying from 2005 onward across the Guiana Shield (Fig 1) (Peek et al. 2005; Carmargo et al. 2009; Breeveld et al. 2012; Evans et al. 2012; Legrand et al. 2012; Musset et al. 2014; Chenet et al. 2017). Worldwide, a handful of validated variants are known to strongly impact parasite phenotype when treated with antimalarial drugs, and several of these variants have documented fitness costs in the absence of drug or collateral sensitivity (*i.e.*, a mutation decreases susceptibility to one drug while increasing susceptibility to another). To determine whether the frequencies of known resistance genotypes changed after widespread ACT adoption, we divided our samples into two groups based on the time of their collection. The first set of 25 genomes was sampled between 1998 and 2005 (“Pre-ACT”); the second set of 21 genomes was sampled between 2013 and 2015 (“Post-ACT”).

**Figure 1.**
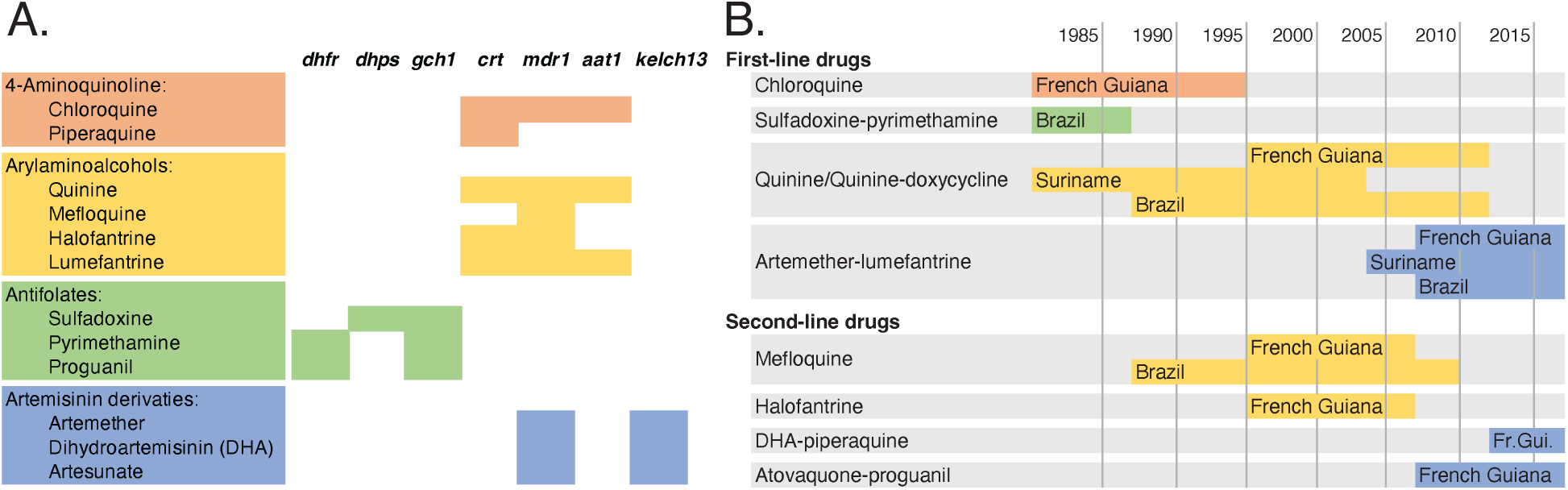
Key genes implicated in resistance to major P. falciparum drug classes (A) and official drug use within French Guiana and its bordering countries (B). While official drug use is depicted here, the region also experienced drug pressure from unregulated sources, especially in gold mining areas. Drugs available on the informal market include: DHA-piperaquine-trimethoprim, Artesunate-sulphamethoxypyrazine-pyrimethamine (Co-Arinate), DHA monotherapy, chloroquine, and primaquine. Key resistance genes are reviewed in Heinberg and Kirkman (2015), Cowell and Winzeler (2019) and Wicht et al. (2020).

We profiled parasite genotypes within six genes harboring known variants that impact resistance to quinolines, antifolates, or artemisinin derivatives and their partner drugs (Table 1, Fig 1). Nearly all alleles show consistent frequencies across the two time periods despite the change in drug pressure and known fitness costs of several alleles. Many, but not all, of these variants have been under genetic surveillance in the Guiana Shield for over two decades, and our results largely recapitulate this previous work (Peek et al. 2005; Legrand et al. 2012).

**Table 1.** SNPs sampled within key resistance-implicated genes.

| Gene | ID | AA position | Allele | Frequency |  |
| --- | --- | --- | --- | --- | --- |
|  |  |  |  | 1998-2005 | 2013-2015 |
| *dhfr* | PF3D7_0417200 | 50 | C50R | 0.96 | 1 |
|  |  | 51 | N51I | 1 | 1 |
|  |  | 59 | C59R | 0.04 | 0 |
|  |  | 108 | S108N | 1 | 1 |
| *mdr1* | PF3D7_0523000 | 184 | Y184F | 0.96 | 1 |
|  |  | 1034 | S1034C | 0.36 | 0.26 |
|  |  |  | S1034C->S | 0.60 | 0.74 |
|  |  | 1042 | N1042D | 0.96 | 1 |
|  |  | 1246 | D1246Y | 0.94 | 1 |
| *aat1* | PF3D7_0629500 | 231 | V231D | 1 | 1 |
|  |  | 258 | S258L | 0.10 | 0 |
|  |  | 446 | P446A | 0.92 | 1 |
| *crt* | PF3D7_0709000 | 72 | C72S | 1 | 1 |
|  |  | 76 | K76T | 1 | 1 |
|  |  | 220 | A220S | 1 | 1 |
|  |  | 326 | N326D | 1 | 1 |
|  |  | 350 | C350R | 0.04 | 0.50 |
|  |  | 356 | I356L | 0.95 | 1 |
| *dhps* | PF3D7_0810800 | 437 | A437G | 1 | 1 |
|  |  | 540 | K540E | 0.94 | 1 |
|  |  | 581 | A581G | 0.96 | 1 |
| *kelch13* | PF3D7_1343700 | 189 | K189T | 0.60 | 0.32 |

One notable exception is a previously unidentified mutation at the second codon position of MDR1^1034^ (PF3D7_0523000). This mutation occurs adjacent to a previously identified high frequency coding mutation, and in combination, the two mutations restore the ancestral amino acid at this position. For over two decades, MDR^S1034C^ (encoded by an A -> T mutation at the first codon position) has been recognized as contributing to multi-drug resistance in South America. Within the Guiana Shield, genetic surveillance approaches have explicitly targeted this single nucleotide polymorphism, sometimes relying on PCR primers that overlap the adjacent second and third codon positions (Peek et al. 2005; Legrand et al. 2012). Here, using whole genome sequencing, we show that the population has in fact returned to having a high frequency of the ancestral amino acid (Ser; 74%) at position 1034, but now encoded by an alternate codon. This change happened before the introduction of ACTs, however, as the MDR^S1034C->S^ variant was already at 60% prevalence in the Pre-ACT time period.

A second exception is *amino acid transporter 1* (*aat1*; PF3D7_0629500), which we profile here for the first time within the Guiana Shield. *In vitro* work recently connected two mutations within this gene to increased chloroquine resistance (Amambua-Ngwa et al. 2023). We find one of these studied variants (S258L) in a single French Guiana parasite sampled in 1999. Along the Pacific Coast of South America, this allele reached near fixation by 2015 (Carrasquilla et al. 2022), but in the Guiana Shield, adaptation appears to have progressed through a pair of alternate mutations: V231D and P446A. This combination of alleles may represent a mode of adaptation unique to the Guiana Shield and Amazon regions of South America; within a set of over 16,000 globally distributed genomes (MalariaGEN et al. 2023), the combination of V231D and P446A is limited to samples from the Guiana Shield and Peruvian Amazon, while V231D alone has been seen in only a handful of samples from West Africa, South Asia, and Southeast Asia (N = 29).

Our remaining results concord with previous Guiana Shield work that has shown the fixation—or near fixation—of known resistance alleles in CRT, MDR1, DHFR, and DHPS across both time periods (Peek et al. 2005; Breeveld et al. 2012; Legrand et al. 2012). An already studied exception is a measurable rise in frequency of the CRT^C350R^ mutation. As with the AAT1 alleles discussed above, CRT^C350R^ is unique to the Guiana Shield and confers piperaquine resistance while reducing chloroquine resistance on the otherwise resistant CRT^K76T^ background (Pelleau et al. 2015; Florimond et al. 2024). We also note a decrease in frequency of the Kelch13^K189T^ variant, although this particular Kelch13 allele is outside the propeller domain and carries no known fitness implications (Amaratunga et al. 2019; Mathieu et al. 2020; Ndwiga et al. 2021)

### Haplotype analysis of temporally spaced samples identifies distinct selective sweeps before and after ACT introduction

In addition to alleles of known effect, we investigated whether the parasite genomes in the Guiana Shield showed further signatures of strong selection after the introduction of ACTs and whether this selection included new genomic targets. As shown in our previous work, mean relatedness between parasites (as measured by identity by descent; IBD), increased across our sampled years (Fig S1A; (Early et al. 2022)). Examining mean pairwise IBD within genomic windows showed that mean relatedness was highly variable across the genome before and after the introduction of ACTs. This pattern is suggestive of selection acting on discrete genomic regions at both time points, however, IBD only provides a measure of population-level relatedness in aggregate. We therefore explored this hypothesis with more rigorous haplotype-based selection tests that analyze frequencies of distinct haplotypes.

We had two main considerations for choosing a haplotype-based summary statistic for this analysis. First, we required sensitivity to both hard and soft sweeps as there is evidence that drug resistance may initially arise via soft sweeps in *P. falciparum* populations (Nair et al. 2007; Anderson et al. 2017). Second, we favored using a simple underlying model given the lack of a fine-scale recombination map for *P. falciparum* and the unusual demographic characteristics of this parasite population (*e.g.*, low and unevenly distributed marker density, recent bottleneck(s), high linkage disequilibrium, and intermittent clonal reproduction). This led us to choose H12 (Garud et al. 2015), which uses predefined haplotype blocks that we could systematically investigate in additional follow-up analyses. H12 is a measurement of haplotype homozygosity that reflects both the number of distinct haplotypes within a genomic window and their relative frequencies. Larger values of H12 arise when a defined window contains one or two “high” frequency haplotype(s), where “high” can be defined in relation to background levels of diversity. As *P. falciparum* harbors much lower diversity and higher linkage disequilibrium than organisms previously studied with H12, the application of the statistic required recalibration of the optimal window size.

As before, we divided our samples into temporally separated sets based on when the use of ACTs became widespread (“Pre-ACT” and “Post-ACT”). Within each sample group, we calculated H12 statistics over a range of window sizes based on either SNP count or physical distance. We chose to define window size in these two different ways as optimal window size is a function of time since selection, local recombination rate, and marker density, all of which vary across the *P. falciparum* genome. We evaluated the performance of the various window sizes with the Pre-ACT data using five previously validated selected loci (chromosome 4: *dhfr*; chromosome 5: *mdr1*; chromosome 6: *aat1*; chromosome 7: *crt*; chromosome 8: *dhps*). With this approach, we found that distance-based windows of 200 kb performed well, identifying four of five known selected loci (Fig 2). In contrast, the SNP-count based approach failed to reflect known sweeps that occurred on chromosomes 4, 5, and 8, likely due to long stretches of extremely low diversity around these regions (Supp Fig S2 and S3). In addition, statistics within SNP-count based windows correlate with the physical size of the window and create what are likely spurious signatures of selection in short, marker-dense windows (Sup Fig S4).

**Figure 2.**
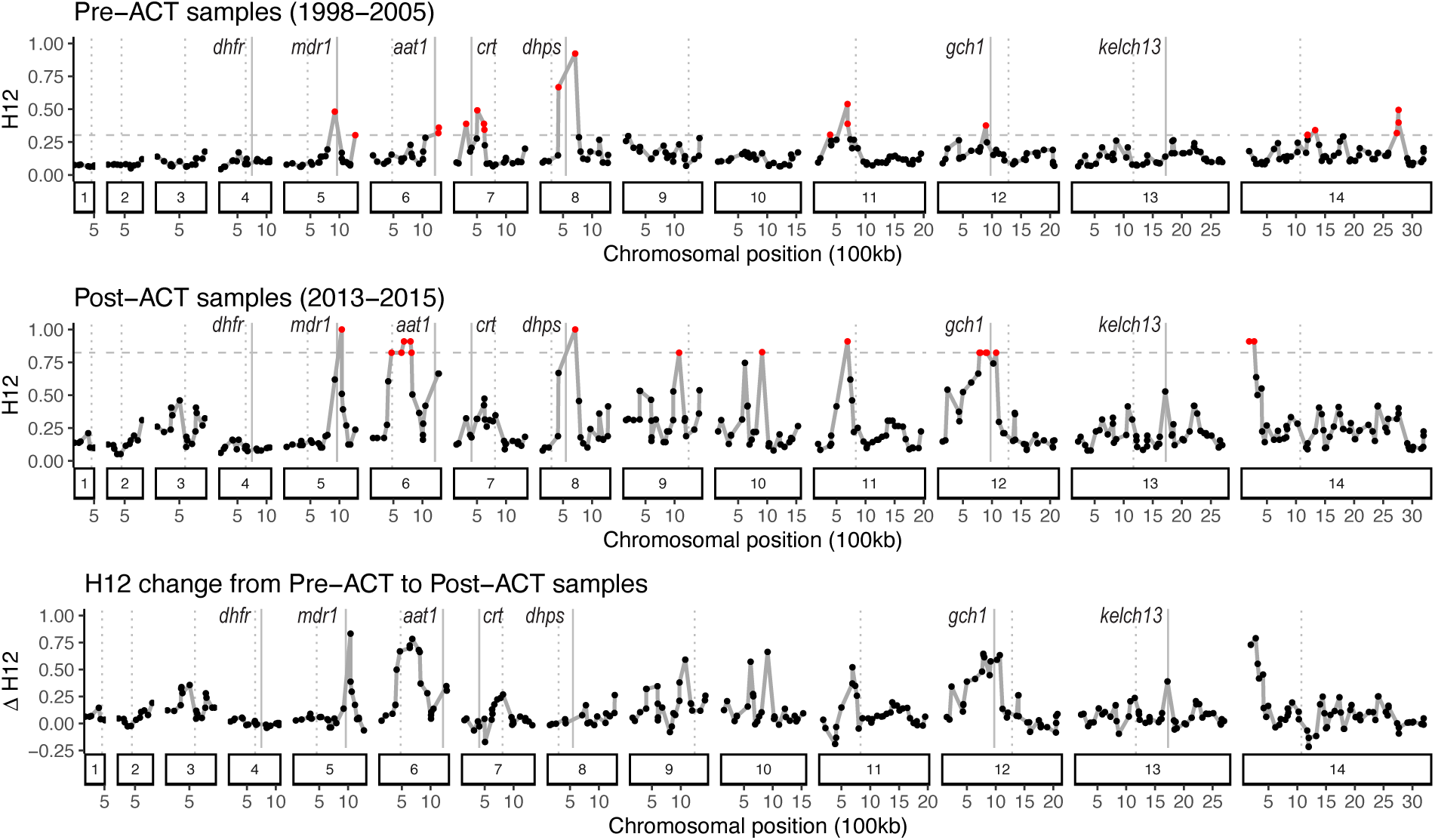
Patterns of haplotype diversity show distinct selection peaks before and after the widespread introduction of ACTs. The H12 statistic was calculated within overlapping 200-kb windows genome wide, with higher values signifying greater haplotype homozygosity. Windows are colored red if they are at or above the 95^th^ percentile for H12 statistics genome-wide (grey dashed line), calculated separately for each time period. Solid vertical lines mark the locations of genes with known drug resistance phenotypes (Chr4: *dhfr*, Chr5: *mdr1*, Chr6: *aat1*, Chr7: *crt*, Chr8: *dhps*, Chr12: *gch1*, Chr13: *kelch13*). Dotted lines mark the location of centromeres. Scans with alternate window sizes are in Supp Figs S2-S5.

In the Pre-ACT sample set, 200-kb windows recovered four H12 peaks that overlap known resistance-associated genes (chromosome 5: *mdr1*; chromosome 6: *aat1*; chromosome 7: *crt*; chromosome 8: *dhps;* Fig 2 and S5). Previous work has shown that these genes experienced strong selection prior to 2000 (Fig 1), and we similarly found mutations within them to be fixed, or nearly fixed, in the Pre-ACT sample set (Table 1). To examine additional peaks, we would ideally determine a significance cutoff through demographic modeling (Harris et al. 2018), however, this approach is currently too inaccurate due to multiple unknowns including the realized recombination rate, changes in effective population size, and population substructure within this relatively small and fragmented population. Instead, we examined the peak size at the four validated markers and noted that all these windows fell within the 95^th^ percentile. Using this as a guidepost, we found four additional chromosomal regions with similarly extreme H12 values (Supplementary Data S1). The genes within these peaks are listed in Supplementary Data S2.

In the Post-ACT sample set, the absolute value of the H12 statistics are higher genome-wide (Fig 2). This reflects the presence of fewer distinct haplotypes in the Post-ACT set relative to the Pre-ACT set. This change is expected due to the increase in average genome-wide relatedness between samples (Fig S1). In general, the Pre-ACT H12 peaks follow this genome-wide trend: they are still present in the Post-ACT set and generally carry higher absolute H12 values. (Although one notable exception is the peak on chromosome 7 that spans the gene *crt*.) This raises a question that we investigated further below: Did selection continue to act at these peaks, or did they drift to higher H12 values as the population contracted?

More conclusively, the Post-ACT analysis shows four new peaks above the 95th percentile, including two multi-window peaks that are present on chromosomes 6 (positions 410,522-912,669; 131 genes) and 14 (positions 40,373-337,115; 79 genes; Supplementary Data S1 & S2). These latter peaks are also seen in the SNP-count based windows (Fig S6), increasing our confidence that the signal at these regions is not due to noise caused by low SNP density. Because of our longitudinal sampling, we can place temporal bounds on when these selection events occurred, dating their start to around the time of ACT introduction. This suggests that nearly complete sweeps occurred in under a decade. The rapidity of these hard sweeps is in keeping with the selection signal being maintained over multiple adjacent windows.

### Most selected regions identified via haplotype analysis show characteristics of hard sweeps

We next investigated the nature of the potential sweeps as there is evidence of both hard and soft sweeps driving *P. falciparum* drug adaptation (Takala-Harrison et al. 2015; Anderson et al. 2017; Amato et al. 2018). To assess the relative likelihood of soft versus hard sweeps at the H12 peaks, we used the H2/H1 statistic which compares haplotype homozygosity after excluding the most common haplotype (H2) to haplotype homozygosity with all haplotypes (H1) (Garud et al. 2015).

For the putative sweeps occurring after 2005, we find hallmarks of hard sweeps. At these genomic windows, H2/H1 approaches zero at the center of the sweep (Fig 3). These low values signal that minor haplotypes contribute little to the overall haplotype homozygosity signal. This phenomenon is similarly summarized by plotting the frequency of the majority haplotype in each window (Fig 4). In the regions of putative selective sweeps on chromosomes 6, 12, and 14, the majority 200-kb haplotype in the 2013-2015 population reached a frequency of 0.95, 0.81, and 0.95, respectively. In these chromosomal regions, single haplotypes therefore rose to near fixation in under a decade despite low, patchy parasite transmission.

**Figure 3.**
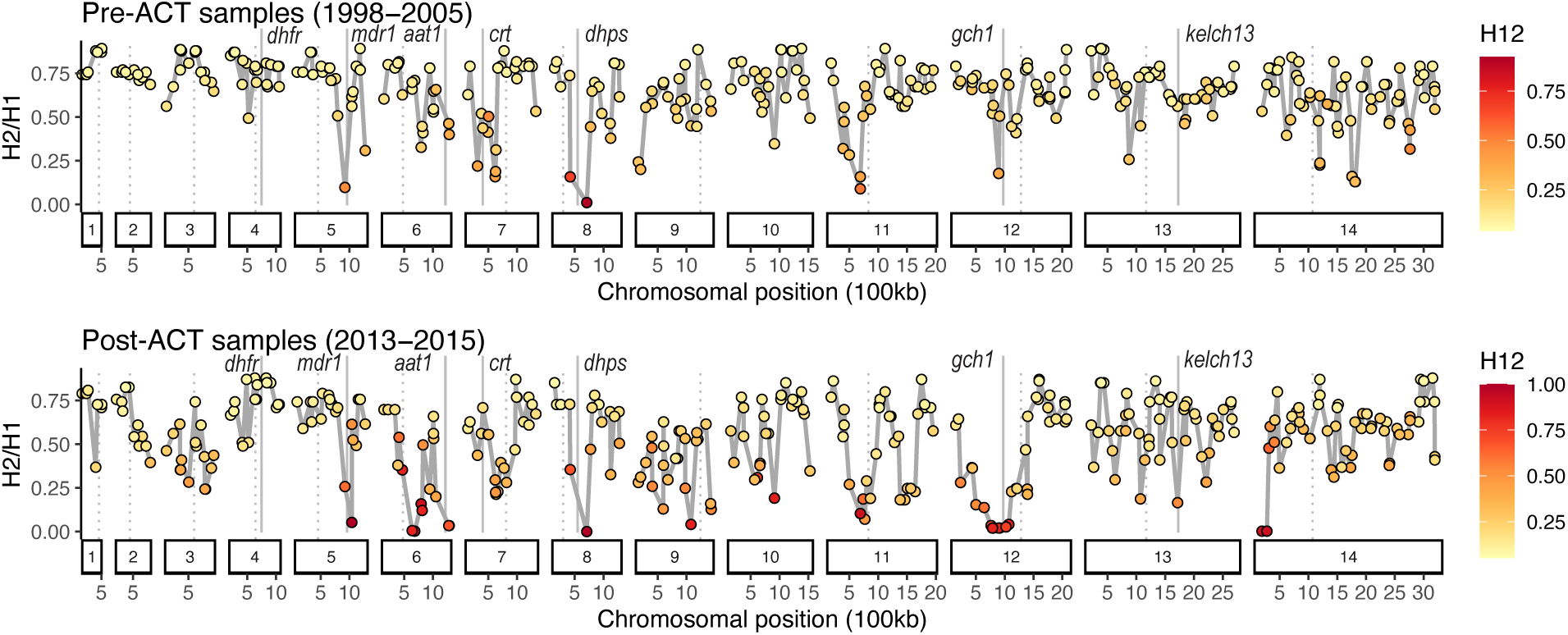
Selection events identified with haplotype analysis are predominated by hard sweep signatures. An H2/H1 value close to 0 corresponds with a model of a hard selective sweep, with values being most informative at the center of putative sweeps (as indicated by high H12 values). For the Post-ACT (2013-2015) group (bottom), most sweeps show evidence of being hard. The H2/H1 values are less extreme in the Pre-ACT group (top), which could indicate either softer sweeps or greater variation in the time since the sweep occurred.

**Figure 4.**
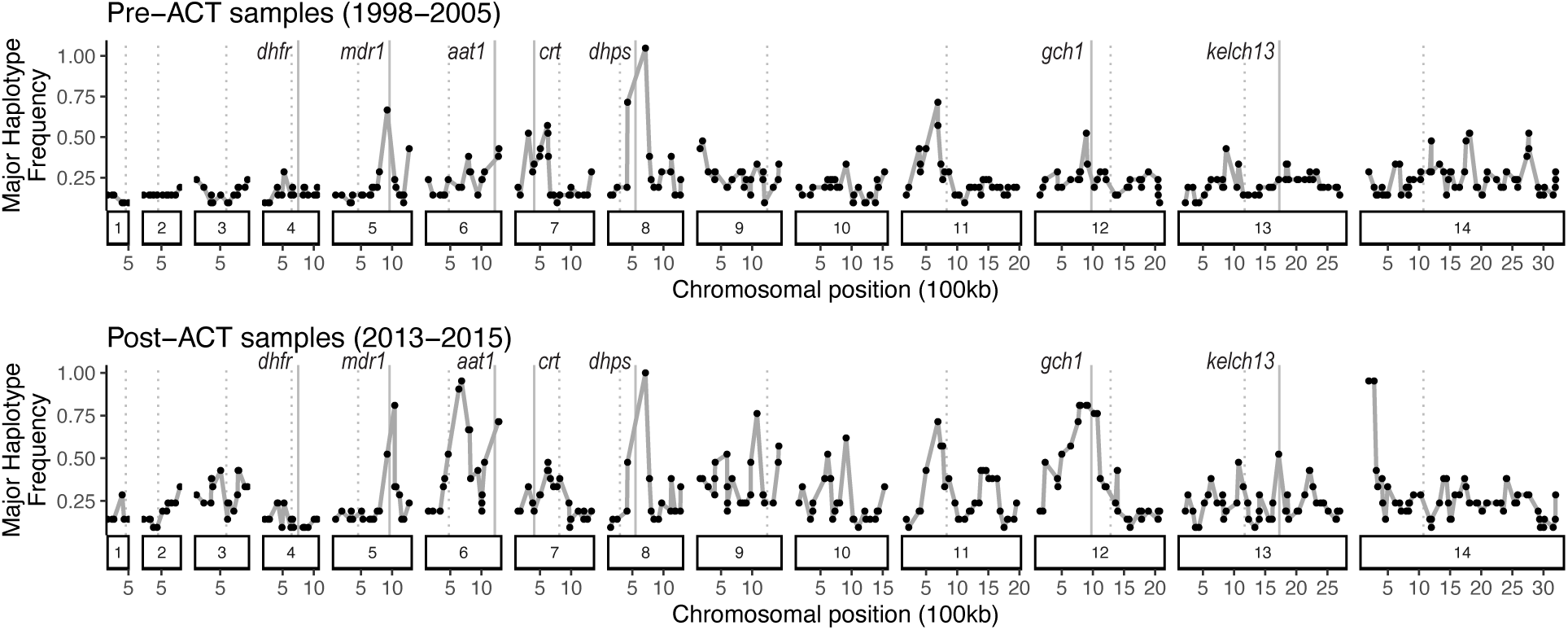
Single haplotypes have approached fixation in novel genomic regions that experienced selection after the introduction of ACTs. Before the introduction of ACTs (top), *P. falciparum* populations contained multiple high frequency haplotypes, largely overlapping loci with known resistance phenotypes. After the widespread introduction of ACTs (bottom), several haplotypes at new genomic targets rapidly approached or reached fixation. The major haplotype was defined independently for each sample set and in some instances, the identity of the major haplotype changed through time (Fig 5).

The H2/H1 values for the Pre-ACT data are less extreme but still quite low at the center of sweeps. This difference might reflect the incomplete nature of some sweeps and/or the greater time over which the Pre-ACT sweeps occurred (which would have allowed recombination or mutation to increase haplotype diversity).

### Multiple genomic regions show consistent selection but a change in favored haplotype after ACT introduction

In general, the H12 peaks present in the Pre-ACT samples are still visible in the Post-ACT samples (Fig 2), and in some instances even appear amplified. This observation concords with the consistent allele frequencies at resistance-conferring loci (Table 1), and suggests at the very least that selection may not have ordinally reversed, and may in some cases have been maintained, following the switch in frontline drug. We, however, investigated the possibility that the temporally maintained H12 signals resulted from sequential bouts of selection on distinct haplotypes at the same locus, rather than consistent selection on a single haplotype. In doing so, we found instances where the major haplotype at a selected locus changed.

In the Pre-ACT data, there are five H12 windows over the 95th percentile that overlap the coding regions of known resistance-conferring genes (Fig 1A): *mdr1* (chr 5), *aat1* (chr 6), *crt* (chr 7), *dhps* (chr 8), and *gch1* (chr 12). At *mdr1*, *crt*, and *dhps*, haplotypes with a high Pre-ACT frequency (>= 0.2) were generally retained at high frequency in the Post-ACT samples (Fig 5 and S7), although we note that an adjacent window overlapping *mdr1* (chr 5)—a window that was not under selection in the Pre-ACT samples—shows a novel signature of positive selection in the Post-ACT samples (Fig 2). A contrasting trend is found around *aat1* and *gch1*; at these two loci, previously favored haplotypes show lower fitness after ACT introduction and were largely replaced by new or previously low frequency haplotypes. Therefore at *mdr1*, *aat1*, and *gch1*, we find evidence of directional selection under both drug regimes, but the selection acted on different haplotypes in the two time periods. Of note, we do not detect any coding variants with frequency changes at these loci. For *gch1* and *mdr1*, selection may be acting on altered expression levels rather than coding changes, as has been seen in this (Legrand et al. 2012) and other populations (Nair et al. 2007; Nair et al. 2008). Unfortunately, we obtained variable sequencing coverage of our samples through time, making copy number assessment from the short-read data unreliable. At *aat1*, we note the possibility that selection may be targeting nearby genes rather than *aat1*. For instance, two genes involved in the *Plasmodium* folate pathway (the target of antifolate drugs; Fig 1) are within the 200-kb region surrounding *aat1*: 6-pyruvoyltetrahydropterin synthase (*ptps*; PF3D7_0628000) and PF3D7_0630700, a putative bifunctional methylenetetrahydrofolate dehydrogenase/cyclohydrolase(Müller and Hyde 2013).

**Figure 5.**
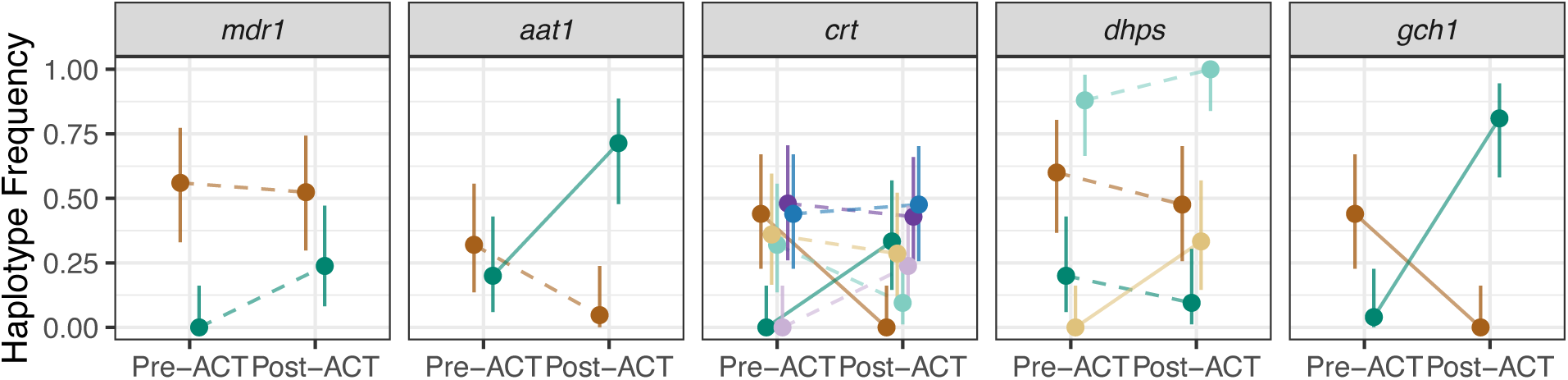
At some consistently selected loci, the favored haplotype changed after the introduction of ACTs. Frequencies are plotted for distinct haplotypes derived from the 200-kb windows that were outliers in the Pre-ACT H12 test (Fig 2) and that overlapped coding regions of known resistance-conferring genes: *mdr1* (chr 5), *aat1* (chr 6), *crt* (chr 7), *dhps* (chr 8), and *gch1* (chr 12). Haplotype frequency shifts sampled from all Pre-ACT H12 outlier windows are in Fig S7. Only haplotypes with a frequency >= 0.2 in at least one time bin are included. Line type represents the significance of the frequency change as determined under a binomial sampling model (dashed: *P* > 0.05, solid: *P* <=0.05). Note that some genes are included in multiple windows, which causes the haplotype frequencies for a gene to have a sum greater than 1. In the case of *mdr1*, two adjacent windows overlapped the gene; only one was an outlier in the Pre-ACT sample set and so is plotted here. The second *mdr1* window was an outlier in the Post-ACT sample set only (Fig 2).

Curiously, at *crt*—the one site of validated selection after 2005 (Pelleau et al. 2015)—we did not see evidence of strong directional selection in the Post-ACT samples (Fig 2). Here again, the identity of the haplotypes themselves is informative. In this instance, we see a loss of the previously favored haplotype that was at a high frequency (0.44) in the Pre-ACT samples but not sampled again after 2005. This lost *crt* haplotype contained the ancestral C350 allele, whereas the haplotypes that increased in frequency after 2005 all carried the derived allele C350R, which experienced strong recent selection due to its impact on chloroquine sensitivity and piperaquine resistance (Table 1).

To place these results in a genome-wide context, we calculated the correlation between the frequency of each Pre-ACT haplotype (n = 6,030 haplotypes) and the same haplotype’s frequency in the Post-ACT samples (Fig S8). These data provide a frame for understanding the stochastic variation present genome-wide due to demographic changes and our relatively small sample sizes. A positive correlation only exists for very common Pre-ACT haplotypes with a frequency of 0.4 or greater (n = 16; Pearson Correlation Coefficient = 0.81). For lower frequency haplotypes, we find no correlation in frequency across time and even observe a substantial negative correlation for haplotypes sampled only 1-2 times in the Pre-ACT group (frequency < 0.1; Pearson Correlation Coefficient = -0.48). Indeed, of the 6,030 Pre-ACT haplotypes defined genome-wide, we observe only 465 (7.7%) in the Post-ACT samples. Overall haplotypic diversity was lower in the Post-ACT samples, with 4,308 haplotypes found genome-wide. Note that while these patterns are consistent with the effects of drift and the stochastic loss of low frequency haplotypes, the magnitude of the trend may also be exacerbated by the smaller sample size in the Post-ACT set (21 vs. 25) and a higher degree of missing genotype calls in the Pre-ACT group, which are not imputed but treated as distinct “variants” by H12 when defining haplotypes.

We found that 67 of our 384 windows contained a haplotype with a frequency change outside expected sampling error (as defined by its 95% binomial confidence interval (CI)). Only seven of these haplotypes were instances where the frequency decreased through time, suggesting lower relative fitness in the later time period. In all seven cases, the haplotypes were unobserved in the Post-ACT samples despite having moderate to high frequency (>=0.36) in the Pre-ACT samples. These include the two haplotypes overlapping *crt* and *gch1* that are discussed above (Fig 5) as well as a region on chromosome 9 (positions 110,142-306,153) and two regions on chromosome 14 (1,650,139 – 1,921,536 and 2,733,317 – 2,887,443). The remaining haplotypes with frequency changes outside their 95% CI all increased in frequency from the Pre-ACT to the Post-ACT time periods. The majority of these haplotypes fall within windows comprising the Post-ACT H12 peaks, either the major peaks on chromosomes 6, 12, and 14 (28 windows) or smaller peaks observable on chromosomes 3, 9, 10, and 11 (22 windows). These results agree with the H12 analysis and provide an additional interpretable layer by foregrounding haplotype identity at selected sites.

### Allele frequency shifts after reversal in selection pressure reveal the complex genetic architecture of drug adaptation

Given the evidence of selection reversal at strongly selected loci, we investigated whether additional genes displayed signatures of opposing selection before and after ACT introduction. To capture the action of soft or incomplete sweeps, we leveraged the temporal sampling design of our data and calculated allele frequencies at biallelic sites across all sampled years. We then modeled allele frequency change through time within two distinct time intervals 1998-2007 and 2007-2015. In 1995, French Guiana stopped official chloroquine use and introduced quinine-doxycycline, mefloquine, and halofantrine. These drugs were then phased out by 2007 in favor of the ACT artemether-lumefantrine. While the timing of these drug changes differed somewhat in neighboring countries (Fig 1), surveys of chloroquine resistance in French Guiana reflected growing susceptibility across this earlier time period, indicating phenotypic change within the parasites (Legrand et al. 2012; Pelleau et al. 2015). We therefore tested whether any alleles showed opposing selection under the quinine-doxycycline/mefloquine/halofantrine regime (1998-2005) and the ACT regime (2007-2015).

We calculated allele frequency trajectories for 26,463 nonsynonymous variants found within 4,197 genes, and of these, we identified 351 nonsynonymous mutations that measurably rose in frequency from 1998 to 2007 and subsequently dropped in frequency from 2007 to 2015 (or vice versa). We placed a relatively low bound on the magnitude of the change, requiring the absolute value of the estimated frequency change to be >0.1. Some detected alleles may therefore have experienced relatively static allele frequencies in one or both time periods. We intersected the 235 genes containing these alleles with genes identified in at least four other functional, genomic, or transcriptomic studies of *P. falciparum* resistance to artemisinin or its partner drugs (358 genes total as summarized in (Oberstaller et al. 2021)) and found 38 overlapping genes (1.9-fold enrichment, Fisher’s Exact Test *P* = 0.00088; Supplementary Data S3). Variants in the 11 genes with the greatest literature support are listed in Table 2 and their yearly frequencies are plotted in Figure S9. Of note, Kelch13 is included among these. However, while multiple mutations in this protein can induce artemisinin tolerance, the only coding variant found in Kelch13 within French Guiana is K189T, which has no known drug phenotype (Amaratunga et al. 2019; Mathieu et al. 2020; Ndwiga et al. 2021). This allele may therefore be linked to a different, causal mutation or itself impact a currently unknown phenotype.

**Table 2.** Genes with allele frequency shifts from 1998-2005 to 2007-2015 that show strongest literature-based evidence of involvement in artemisinin or partner drug resistance.

| Gene | ID | Allele |
| --- | --- | --- |
| SURFIN 1.2 (pseudogene) | PF3D7_0113600 | A181E |
|  |  | Q526P |
|  |  | Q536R |
|  |  | G537S |
|  |  | N559D |
|  |  | N939D |
|  |  | Y1015N |
| Plasmodium exported protein | PF3D7_0701900 | S895N |
|  |  | E890Q |
|  |  | T866K |
|  |  | R344S |
| conserved Plasmodium protein | PF3D7_0710200 | Y2240F<br>N2242K |
| conserved Plasmodium membrane protein | PF3D7_0804500 | A4860D |
| HECT-like E3 ubiquitin ligase | PF3D7_0826100 | I380T |
| conserved Plasmodium protein | PF3D7_0922800 | I1000M |
| heat shock protein 90 | PF3D7_1118200 | F475Y<br>H474D |
| autophagy-related protein 7 | PF3D7_1126100 | E979G |
|  |  | Y812C |
| myosin F | PF3D7_1329100 | D348Y<br>N277S |
| kelch13 | PF3D7_1343700 | K189T |
| GTP-binding protein | PF3D7_1462300 | D1339N |

## DISCUSSION

As policy makers design recommendations for drug combinations or drug cycling, it is imperative to consider that drug-induced selection pressures are not fully independent or distinct. This study captures how selection under sequential drug regimes repeatedly targets the same genomic regions in *P. falciparum*, which is consistent with drug changes either unmasking fitness costs or revealing collateral sensitivity in the parasite. By documenting these selection patterns genome wide, our results show that such trade-offs extend beyond the handful of large effect genes that predominate the *P. falciparum* literature. Interestingly, some observed genomic patterns deviate from standard expectations. *A priori*, we would expect to see evidence for relaxed selection when drug pressure is removed, or, if the resistance allele carries a fitness cost in the absence of drug, a reversion to the wildtype allele. In keeping with this model, we find individual alleles whose frequency trajectory changed signs after the change to ACTs (Table 2). Beyond this, however, we document examples where distinct haplotypes at the same locus (*eg, aat1* and *gch1)* were targets of strong directional selection under the different drug regimes (Fig 5). The newly selected haplotypes may have conferred resistance to the new drugs or mitigated the cost of previously fixed alleles. As an example of the latter, the relatively recent MDR1 allele that restores the wildtype amino acid at position 1034 highlights how *Plasmodium* adaptation may occur through novel mutations rather than a reversion to an ancestral nucleotide allele. Overall, these observations are not unlike what has been observed in bacteria where evolutionary trajectories depend on environment, epistatic interactions, and mutation rate (Pennings et al. 2022).

In addition to continued selection at previous targets, we also detect hard sweeps at novel loci that occurred after the introduction of ACTs. These new sweeps were extremely rapid and nearly fixed previously low frequency segments on chromosomes 6 and 14 in under eight years. This shows that, even under strong control measures, both gene flow and effective recombination remained sufficiently high to passage selected haplotypes throughout the whole population. Together with the observation of more subtle allele frequency shifts, these loci point to the likely polygenic nature of drug adaptation and show that small populations harbor sufficient genotypic and phenotypic diversity to adapt via both new mutation and standing variation.

### Connecting genotype to phenotype

Interestingly, the strong selection signatures in the post-ACT samples did not coincide with ACT failure in the Guiana Shield, although initial signs of partner-drug resistance have emerged in the region. Recently, we reported the first evidence of resistance to the artemisinin partner drug piperaquine, which is intermittently used in French Guiana and available on the informal market throughout the Guiana Shield (Florimond et al. 2024). The study found that the mutation CRT^C350R^ partially explains the phenotype and that copy number variation at *plasmepsin II* and *plasmepsin III* potentiates piperaquine resistance when on a CRT^C350R^ background (Florimond et al. 2024). These plasmepsin genes are located in the region of chromosome 14 that showed evidence of strong selection in the Post-ACT samples. Interestingly, copy number at this locus is not fixed in the post-ACT population despite the SNP-based evidence that the same chromosomal segment is shared across all extant parasites. Copy number at the locus therefore shows evidence of being highly dynamic, leaving open the possibility of a hard sweep selecting for a *plasmepsin II/III* duplication followed by copy loss in some descendent lineages (Vanhove et al. 2024).

It is also probable that phenotypes other than drug resistance are under selection. For example, each of the Post-ACT peaks on chromosomes 6, 12, and 14 contains AP2 transcription factors that are key for sexual commitment and gametocyte development (Supplementary Data S2). Both we and others have found evidence that optimal sexual investment rate for *Plasmodium* varies across transmission settings (Rono et al. 2018; Early et al. 2022), which supports these genes as potential selection targets. Unfortunately, the chromosomal regions that experienced strong selection are large, making the exact target of selection impossible to pinpoint (Supplementary Data S1 & S2). Regardless of the ultimate drivers of these sweeps, however, the absence of clinical failure throughout the region highlights the fact that additional phenotypes carry sufficiently large fitness coefficients to drive strong selection. Further work is required to test whether these involve subtle shifts in tolerance or resistance, alternate traits like sexual conversion rate, or compensatory mutations mitigating the cost of previously fixed alleles.

### Adaptation through both common and novel genetic mechanisms

In South America and Southeast Asia, parasite populations have independently evolved identical alleles in key genes while also acquiring population-specific mutations (Anderson and Roper 2005). Here, we identify geographically limited mutations in resistance-relevant genes (MDR1^S1034C->S^, AAT1^V231D^ and AAT1^P446A^) and novel haplotype selection peaks that likely signal region-specific modes of adaptation. The MDR1^S1034C->S^ variant that restored the ancestral amino acid is of particular interest as it evaded detection for over a decade despite genomic surveillance of this locus. Periodic region-specific genome-wide selection scans are therefore warranted to stay abreast of the ever-changing parasite genome.

However, adaptations that are shared between populations are also brought to light through region-specific selection scans. Genomic analyses—whether association tests, expression analysis, or tests of selection—always struggle with separating signal from noise; typically, only the strongest contenders garner downstream attention. As we show here (Table 2), signal enrichment across distinct populations is therefore a fruitful way to identify additional globally relevant candidate genes and unpack the polygenic architecture of *Plasmodium* drug resistance.

### Challenges of selection detection in small populations

While our approach successfully identified genomic regions under selection, the results also highlight current genomic blind spots and the need for further methods development. For instance, nucleotide diversity is unevenly distributed across the genome, a pattern that is further exacerbated when genotyping is based on short-read sequence data. In the H12 analysis, the region around *dhps* on chromosome 8 is striking due to its long peak when using distance-based windows. Without *a priori* interest in the region, however, these windows might have been discarded due to their exceptionally low marker density (Supplementary Data S1), and they show no signal when SNP-based windows were used (Fig S2). Conversely, we only detect a signal in the vicinity of *dhfr* (chromosome 4; Fig S2) when using windows based on 20 SNPs. Further assessing demographic characteristics of this parasite population and subsequently developing appropriate null models will bolster our ability to differentiate the effects of selection versus drift. As of now, we cannot discount that demographic effects might have contributed to the large novel H12 peaks observed in the post-ACT samples. In sum, our ability to discern robust patterns in the data rested heavily on previously acquired knowledge. Similarly, our ability to detect soft and incomplete selective sweeps relied on the longitudinal nature of our data set. Further methodological advances will need to be made before conducting comparable analyses in previously unstudied populations, and investigators should continue to develop longitudinal data sets.

Nevertheless, our ability to reliably detect loci under selection with small sample sizes within a low transmission population is heartening as effective evolutionary analysis in these geographic regions will be essential for preventing malaria adaptation and resurgence. Drift is expected to obscure selection signals in small populations nearing elimination, and indeed our selection tests do contain high levels of background noise. Yet despite this, we detect strong signals and identify not only known but also credible, new resistance markers. Therefore, if incorporated into standard monitoring activity, selection scans like these can be used to guide future public health measures by detecting evidence of successful parasite adaptation before clinical resistance emerges.

## METHODS

### Sample collection, sequencing, and variant calling

French Guiana samples were collected and sequenced as previously described in Early & Camponovo, *et al* (Early et al. 2022). In brief, 148 anonymized blood samples were collected from symptomatic *P. falciparum*-positive cases between 1998 and 2015. We culture adapted forty-three of the isolates prior to sequencing, which resulted in enhanced coverage (Supplementary Table S1). All samples underwent library construction using the Nextera XT low-input library kit followed by whole-genome shotgun on an Illumina HiSeq 2500 to generate 100-bp paired-end reads. We aligned raw reads with BWA-mem to the PlasmoDB *P. falciparum* 3D7 v.2.8 reference genome and called variants with GATK v.3.5 HaplotypeCaller using a set of genotyped pedigreed crosses for base and variant score recalibration. We limited downstream analysis to SNPs called in the “core” genome as defined by (Miles et al. 2016). Raw sequence data are available in the National Center for Biotechnology Information (NCBI) Sequence Read Archive as BioProject PRJNA3361. We calculated relatedness (as identity-by-descent) between all pairs of putatively monogenomic samples using the program hmmIBD (Schaffner et al. 2018). As input, we used allele frequencies calculated across the entire set of samples. At the group level, we summarized IBD as the mean of all pairwise comparisons genome-wide (fractional IBD) or within 100-kb windows.

### Haplotype-based selection analysis

For haplotype-based analysis, we selected 46 monogenomic samples with >=30% of the genome at 5x sequencing depth as identified in (Early et al. 2022). All genomes were separated by at least one recombination event (pairwise fractional IBD < 0.9). We divided samples into time periods before and after the widespread use of ACTs in French Guiana in 2007 (“Pre-ACT”: 1998-2005 N=25; “Post-ACT”: 2013-2015 N=21). To maximize the accuracy of haplotype identification, we rigorously filtered the variants present in these genomes. We filtered out sites if they were (1) called in <80% of samples; (2) within “non-core” regions of the genome; (3) within a hypervariable gene family (*var*, *stevor*, *rifin*); or (4) within a gene showing an unusually high proportion of variant sites (>= 0.005; N=61). We masked all heterozygous calls within individual samples. To enable comparison across sample groups, we used the identical list of sites for both sample sets. As a result, some sites were variant in one group and invariant in the other.

We initially performed selection testing using the package SelectionHapStats (Garud et al. 2015) to calculate H12 and H2/H1 with a range of SNP-based windows. We subsequently applied the statistic to distance-based windows, which the program does not directly support. To do this, we defined coordinate-based windows, identified the SNPs present within the window, and passed the SNP information on to the H12 algorithm. In practice, the realized length of the distance-based windows varied due to variability of marker density across the genome. We therefore calculated the length between the first and last marker within each window and took this metric into consideration when interpreting outlier windows. Due to the low number of windows per chromosome, we did not standardize scores by chromosome and recognize this may create some detection bias in the data.

### Allele frequency changes

We estimated yearly allele frequencies with a pooled approach that included both low-coverage and high-coverage samples. We downsampled all raw BAM files to approximately 1x mean genome coverage and then merged downsampled BAMs from samples collected in the same year. Using the pooled BAMs, we called variants using GATK v.3 and used the read ratios at each variant to estimate yearly allele frequencies.

To calculate allele frequency trajectories at biallelic sites, we performed separate logistic regressions with the estimated allele frequencies at each variant within two time periods: 1998—2007 and 2007—2015. We used the confint function in R to calculate the 95% confidence interval for each estimated slope. To retain the results for a given locus, we required that the models for the two time periods had: (1) at least three (dof=1) observed allele frequency measurements each, (2) estimated slopes with 95% confidence intervals that did not intersect zero, and (3) slopes in opposing directions. This identified 351 nonsynonymous mutations in 235 genes. We additionally analyzed the data with a more stringent filter on missing data, requiring at least four observed allele frequency measurements per time point. This reduced the number of significant hits to 317 nonsynonymous sites in 207 genes, but the qualitative trends remained comparable.

## Notes

### Competing Interest Statement

The authors have declared no competing interest.

