## Supplementary Figures for "Temporal patterns of haplotypic and allelic diversity reflect the changing selection landscape of the malaria parasite *Plasmodium falciparum*"

**A.**

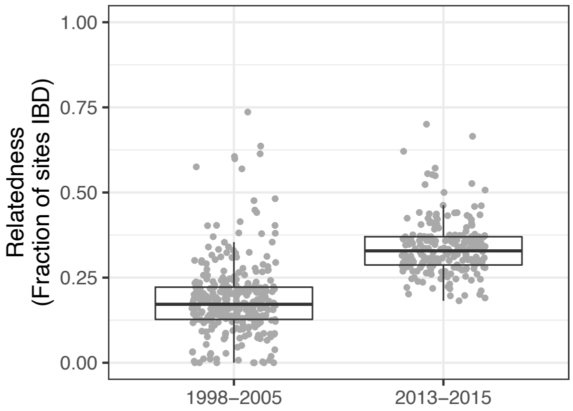


**B.**

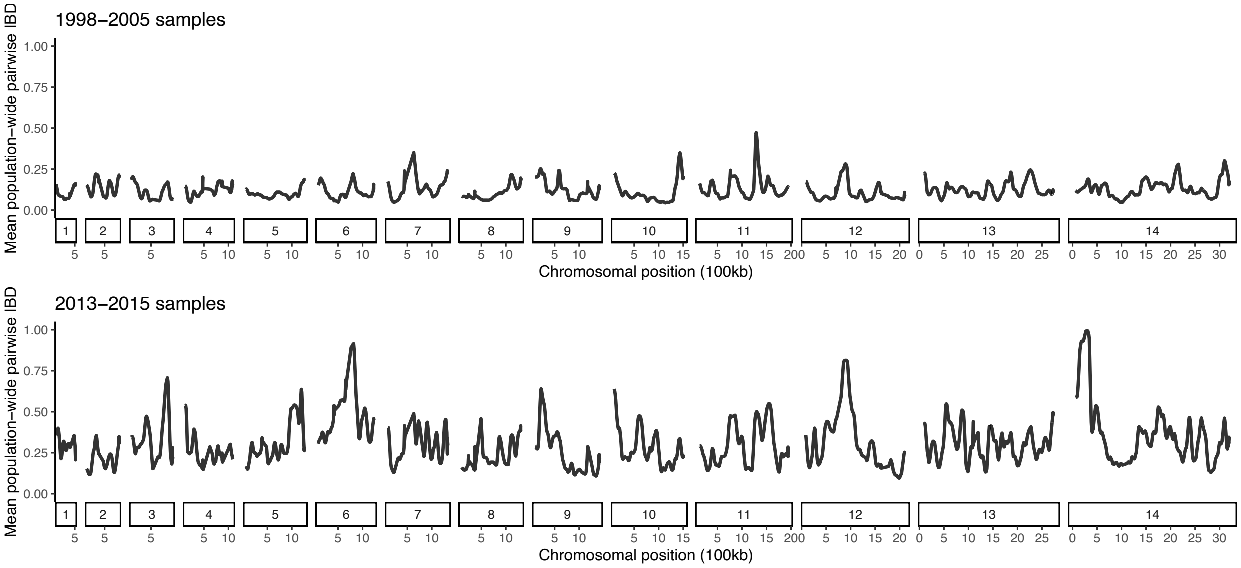


**Figure S1. Relatedness between parasites increased after a successful malaria intervention campaign that began in 2007.** (A) Median fractional IBD between parasites increased from 0.19 in the Pre-ACT group (1998-2005) to 0.31 in the Post-ACT group (2013-2015). (B) Genome sharing increased unevenly across chromosomal regions.

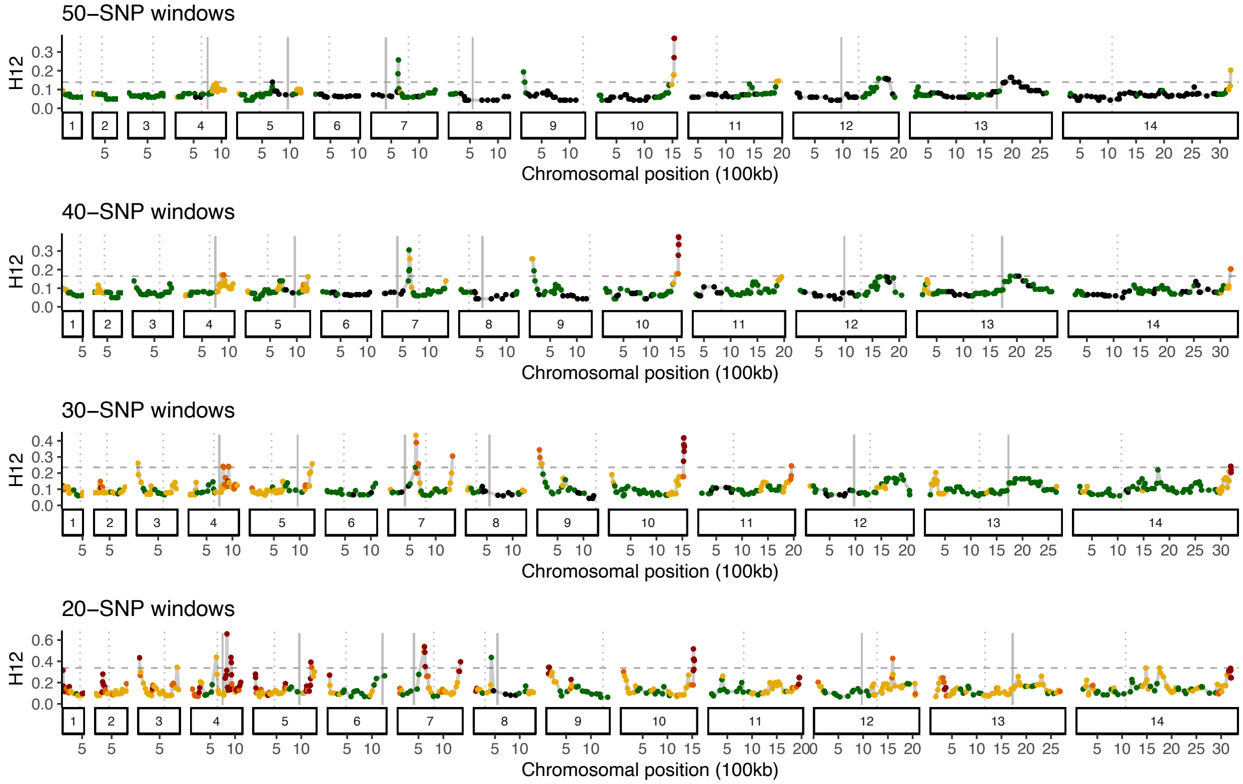


**Figure S2. H12 scans of the Pre-ACT group (1998-2005) across a range of SNP-count based window sizes.** Based on previous work in the population, peaks were expected on chromosomes 4, 5, and 7 (*dhfr*, *mdr1*, and *crt*, respectively), but no window size recovered all of these. Points are colored by physical window size (Red: <75kb, Orange: 75kb-100kb, Red: 100kb-200kb, Green: 200kb-400kb, Black: over 400kb). For reference, the horizontal dashed line denotes the 95$^{th}$ percentile for H12 statics genome-wide. Solid vertical lines mark the locations of genes with known drug resistance phenotypes (Chr4: *dhfr*, Chr5: *mdr1*, Chr6: *aat1*, Chr7: *crt*, Chr8: *dhps*, Chr12: *gch1*, Chr13: *kelch13*). Dotted vertical lines mark the location of centromeres.

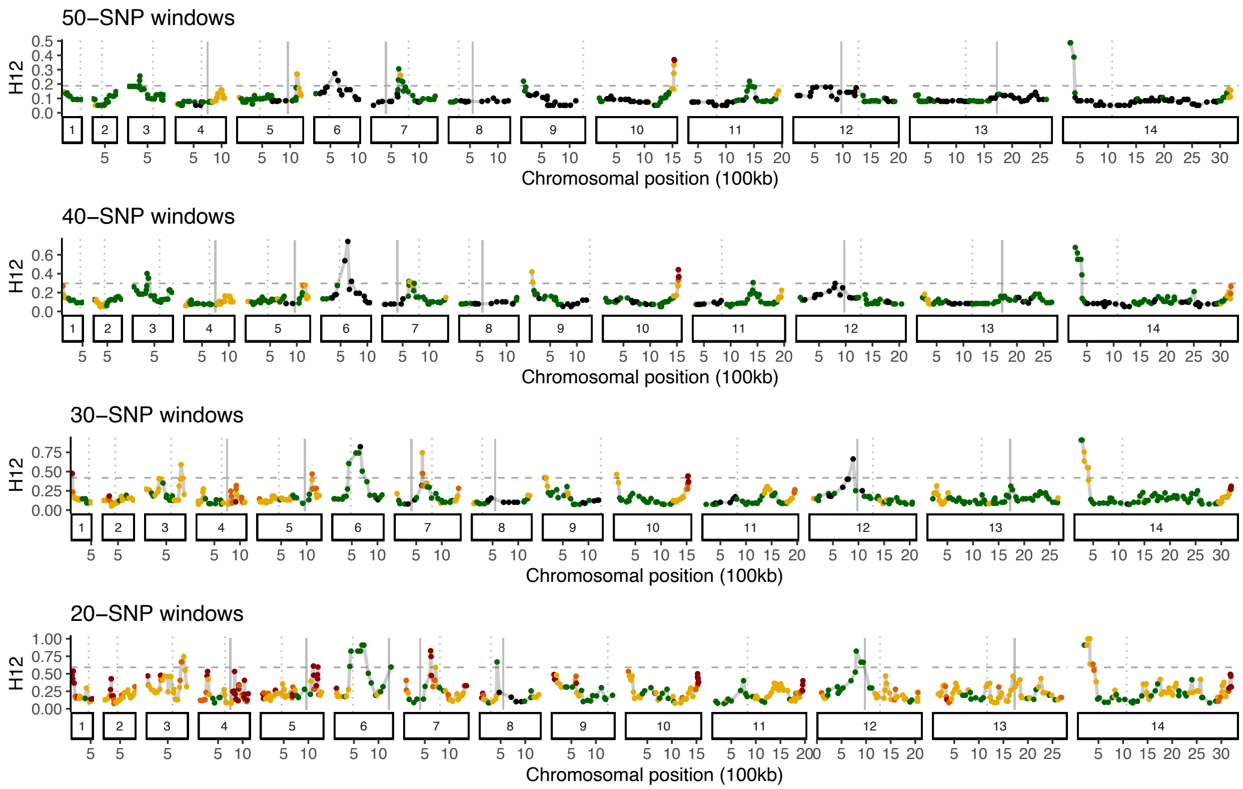


**Figure S3. H12 scans of the Post-ACT group (2013-2015) across a range of SNP-count based window sizes.** Peaks on chromosomes 6 and 14 are the strongest candidates of selection as they are detected across all SNP-count based window sizes and also observed when using distance-based windows. Points are colored by physical window size (Red: <75kb, Orange: 75kb-100kb, Red: 100kb-200kb, Green: 200kb-400kb, Black: over 400kb). For reference, the horizontal dashed line denotes the 95$^{th}$ percentile for H12 statics genome-wide. Solid vertical lines mark the locations of genes with known drug resistance phenotypes (Chr4: *dhfr*, Chr5: *mdr1*, Chr6: *aat1*, Chr7: *crt*, Chr8: *dhps*, Chr12: *gch1*, Chr13: *kelch13*). Dotted vertical lines mark the location of centromeres.

**A.**

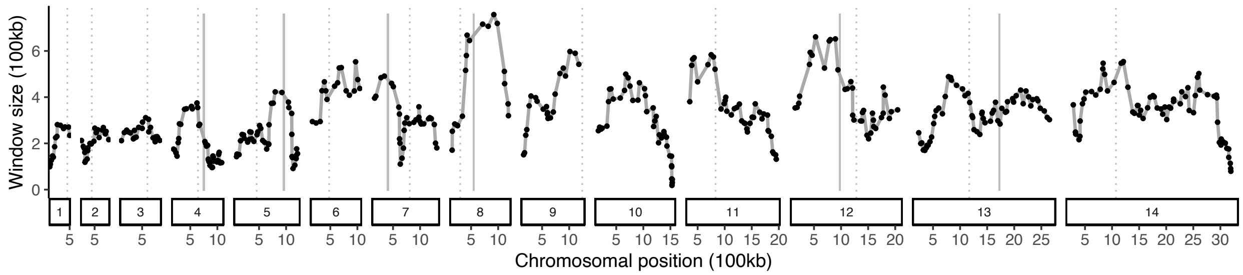


**B.**

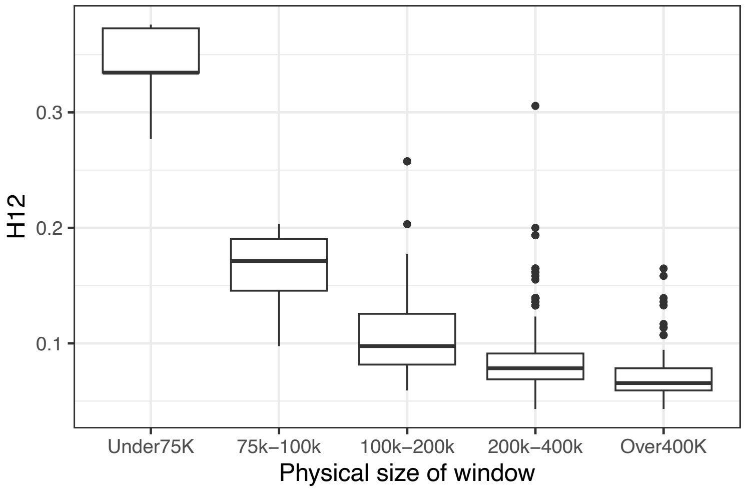


**Figure S4. Variation in the physical size of the H12 windows when defined based on SNP counts**. Windows with a small physical size are (A) enriched at chromosome ends and (B) show inflated H12 values. Data shown are for 40-SNP H12 windows calculated with the Pre-ACT samples. Sub-telomeric regions and genes with a high proportion of variant sites (>= 0.005) were previously removed from the data sets. In (A), Solid vertical lines mark the locations of genes with known drug resistance phenotypes (Chr4: *dhfr*, Chr5: *mdr1*, Chr6: *aat1*, Chr7: *crt*, Chr8: *dhps*, Chr12: *gch1*, Chr13: *kelch13*). Dotted vertical lines mark the location of centromeres.

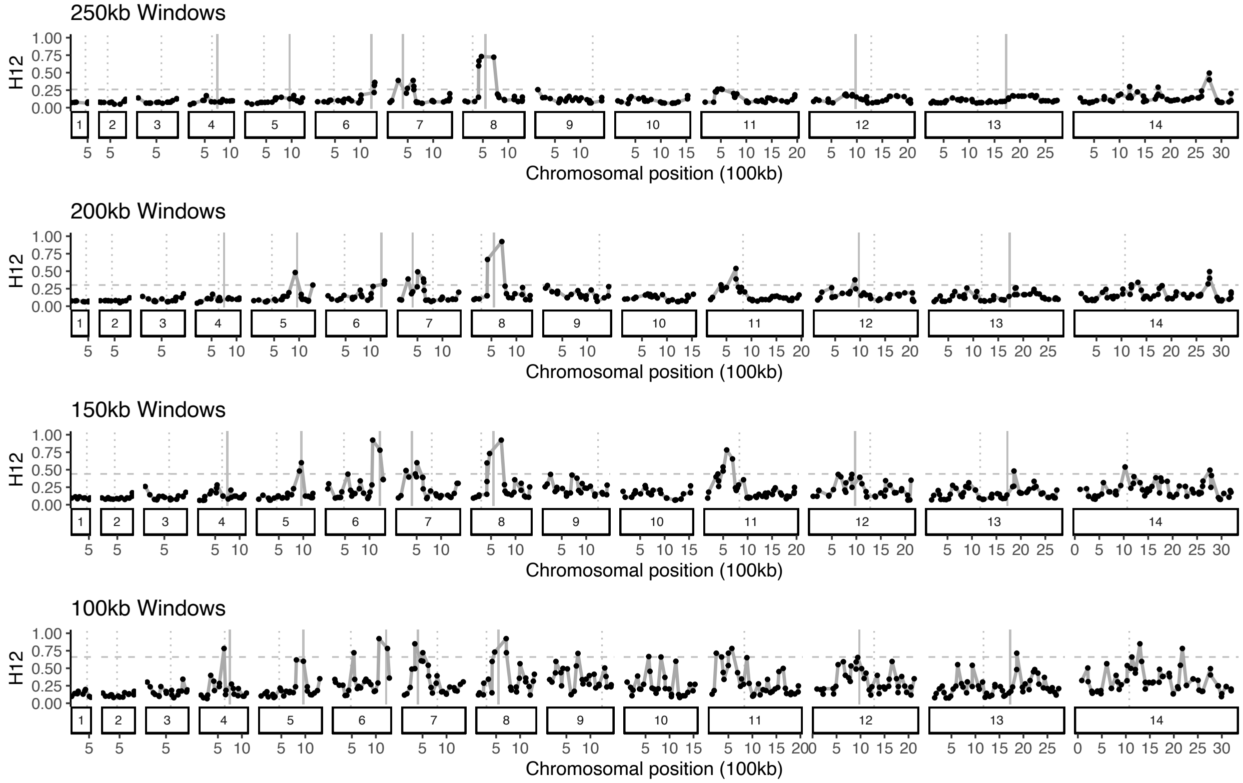


**Figure S5. H12 scans of the Pre-ACT group (1998-2005) across a range of distance-based window sizes.** For reference, the horizontal dashed line denotes the genome-wide 95th percentile. Solid vertical lines mark the locations of genes with known drug resistance phenotypes (Chr4: *dhfr*, Chr5: *mdr1*, Chr6: *aat1*, Chr7: *crt*, Chr8: *dhps*, Chr12: *gch1*, Chr13: *kelch13*). Dotted vertical lines mark the location of centromeres.

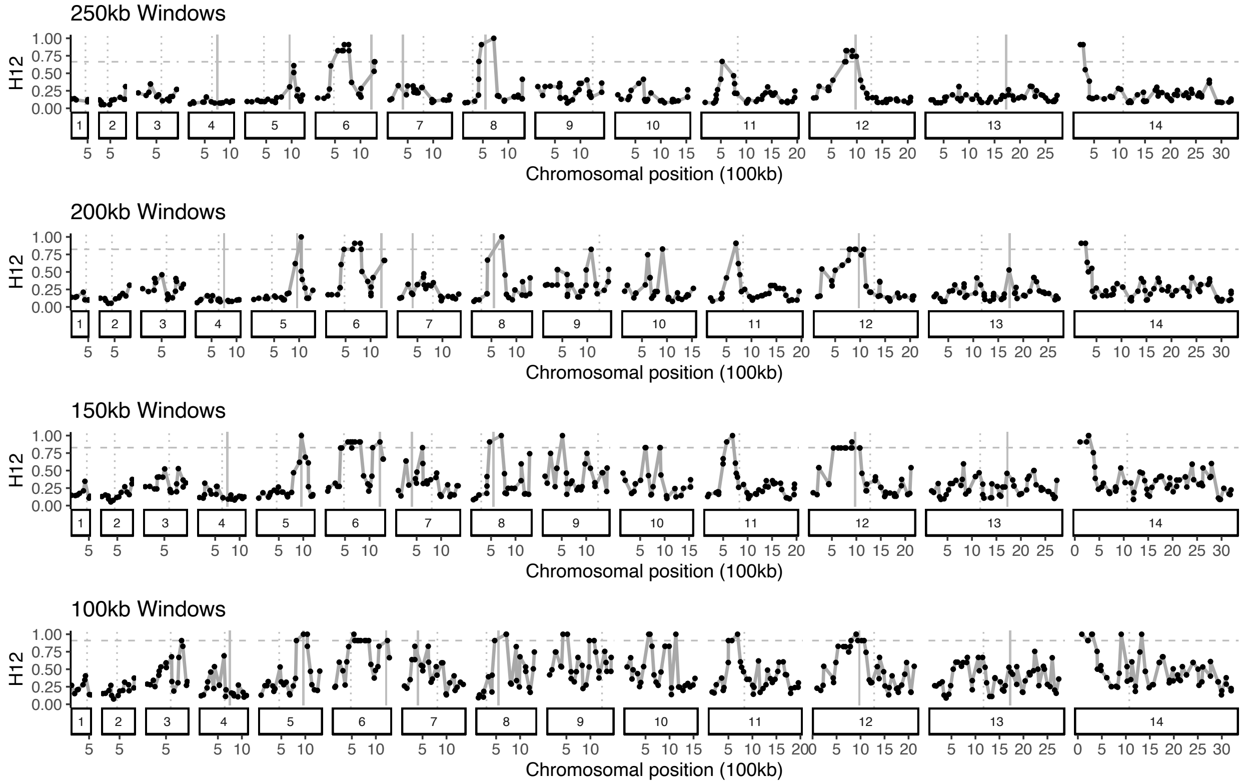


**Figure S6. H12 scans of the Post-ACT group (2013-2015) across a range of distance-based window sizes.** For reference, the horizontal dashed line denotes the genome-wide 95th percentile. Solid vertical lines mark the locations of genes with known drug resistance phenotypes (Chr4: *dhfr*, Chr5: *mdr1*, Chr6: *aat1*, Chr7: *crt*, Chr8: *dhps*, Chr12: *gch1*, Chr13: *kelch13*). Dotted vertical lines mark the location of centromeres.

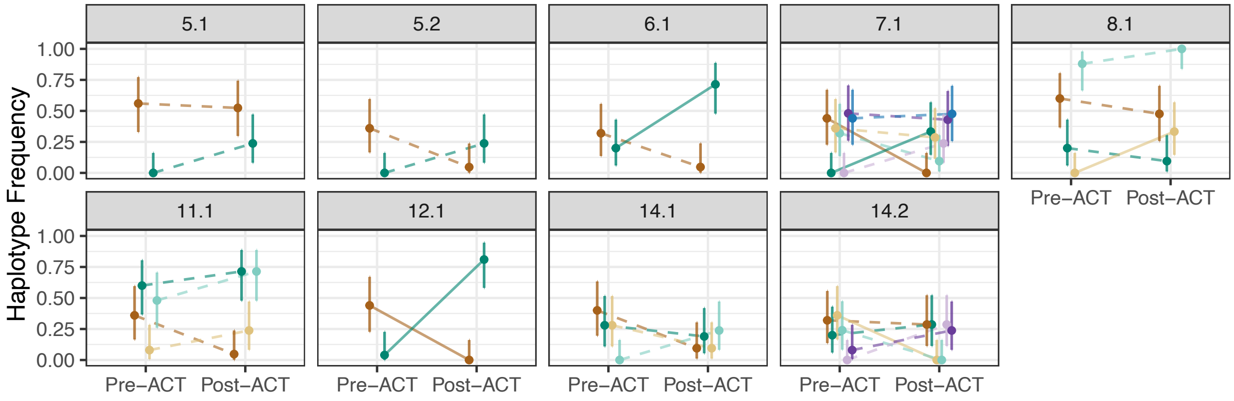


**Figure S7.** Frequencies of all haplotypes from the 200-kb windows that were outliers in the Pre-ACT H12 test (Fig 2) and at a frequency >= 0.2 in at least one time bin. Line type represents the significance of the frequency change as determined under a binomial sampling model (dashed: $P > 0.05$, solid: $P <= 0.05$). All haplotypes from consecutive windows forming a single peak are included in the same subplot.

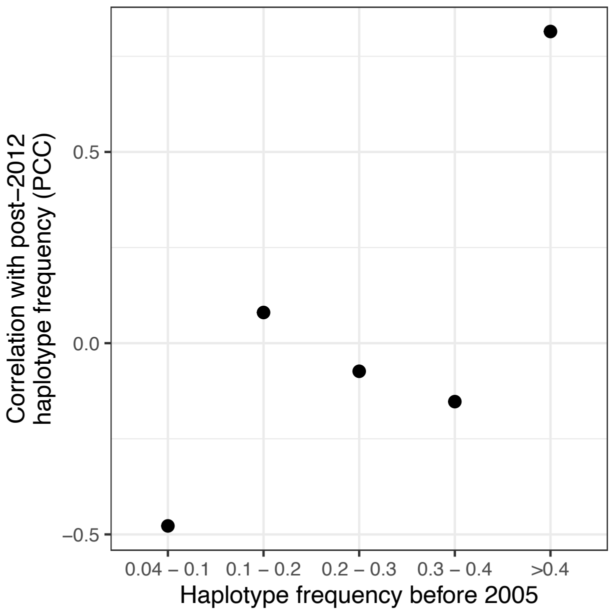


**Figure S8. Correlation between haplotype frequency in the Pre-ACT (1998-2005) and Post-ACT (2013-2015) samples.** The Pearson Correlation Coefficient (PCC) between Pre-ACT and Post-ACT frequency was calculated for all haplotypes within the given frequency intervals in the Pre-ACT samples.

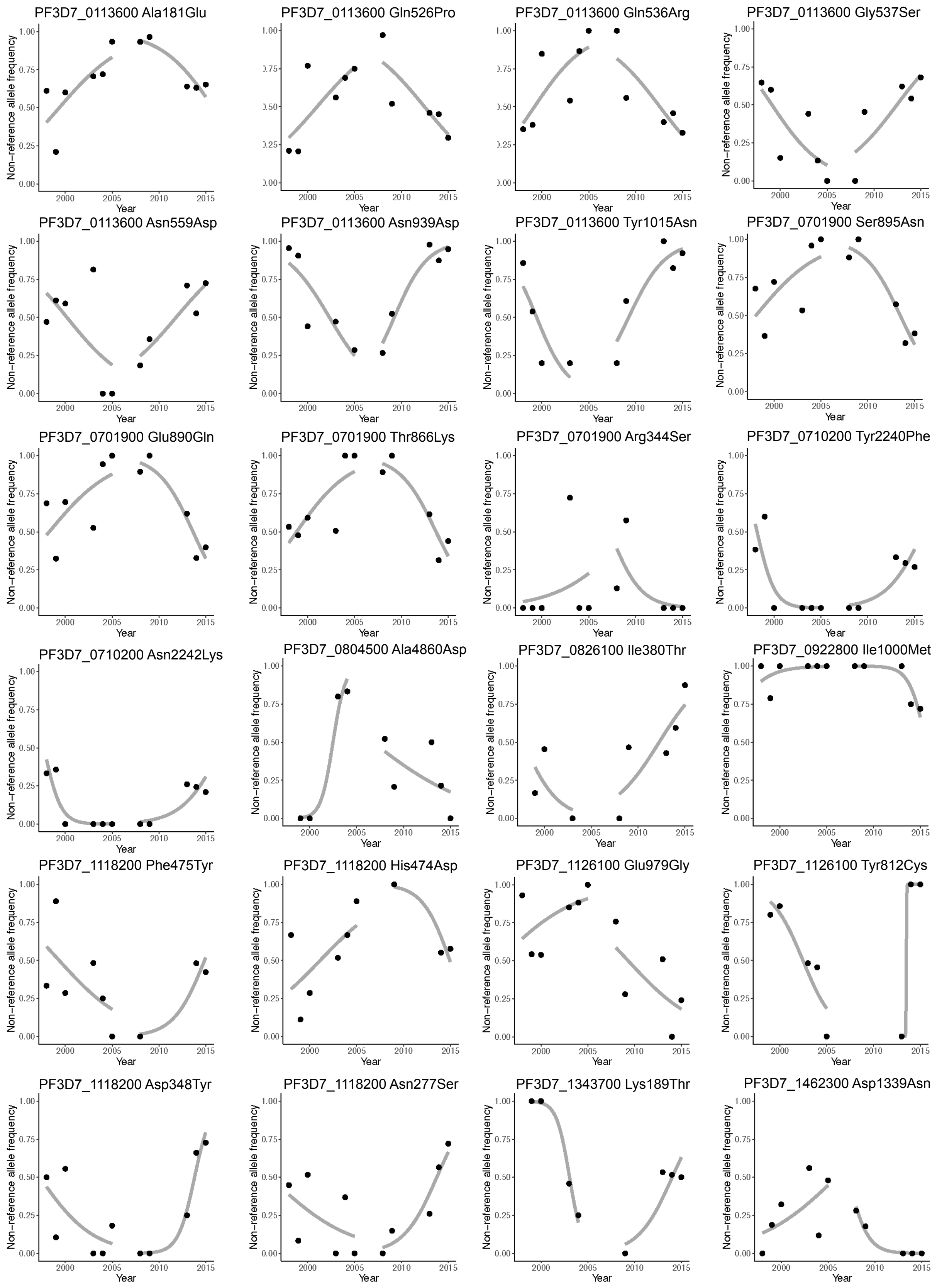


**Figure S9. Allele frequencies through time for the variants listed in Table 2.**
